# TREM2+ and interstitial macrophages orchestrate airway inflammation in SARS-CoV-2 infection in rhesus macaques

**DOI:** 10.1101/2021.10.05.463212

**Authors:** Amit A. Upadhyay, Timothy N. Hoang, Maria Pino, Arun K. Boddapati, Elise G. Viox, Michelle Y.H. Lee, Jacqueline Corry, Zachary Strongin, David A. Cowan, Elizabeth N. Beagle, Tristan R. Horton, Sydney Hamilton, Hadj Aoued, Justin L. Harper, Kevin Nguyen, Kathryn L. Pellegrini, Gregory K. Tharp, Anne Piantadosi, Rebecca D. Levit, Rama R. Amara, Simon M. Barratt-Boyes, Susan P. Ribeiro, Rafick P. Sekaly, Thomas H. Vanderford, Raymond F. Schinazi, Mirko Paiardini, Steven E. Bosinger

**Author notes:** Correspondence to Steven E. Bosinger; Mirko Paiardini. These authors contributed equally (Amit A. Upadhyay, Timothy N. Hoang, Maria Pino).

## Abstract

The COVID-19 pandemic remains a global health crisis, yet, the immunopathological mechanisms driving the development of severe disease remain poorly defined. Here, we utilize a rhesus macaque (RM) model of SARS-CoV-2 infection to delineate perturbations in the innate immune system during acute infection using an integrated systems analysis. We found that SARS-CoV-2 initiated a rapid infiltration (two days post infection) of plasmacytoid dendritic cells into the lower airway, commensurate with IFNA production, natural killer cell activation, and induction of interferon-stimulated genes. At this early interval, we also observed a significant increase of blood CD14-CD16+ monocytes. To dissect the contribution of lung myeloid subsets to airway inflammation, we generated a novel compendium of RM-specific lung macrophage gene expression using a combination of sc-RNA-Seq data and bulk RNA-Seq of purified populations under steady state conditions. Using these tools, we generated a longitudinal sc-RNA-seq dataset of airway cells in SARS-CoV-2-infected RMs. We identified that SARS-CoV-2 infection elicited a rapid recruitment of two subsets of macrophages into the airway: a C206+MRC1-population resembling murine interstitial macrophages, and a TREM2+ population consistent with CCR2+ infiltrating monocytes, into the alveolar space. These subsets were the predominant source of inflammatory cytokines, accounting for ~75% of IL6 and TNF production, and >90% of IL10 production, whereas the contribution of CD206+MRC+ alveolar macrophages was significantly lower. Treatment of SARS-CoV-2 infected RMs with baricitinib (Olumiant^®^), a novel JAK1/2 inhibitor that recently received Emergency Use Authorization for the treatment of hospitalized COVID-19 patients, was remarkably effective in eliminating the influx of infiltrating, non-alveolar macrophages in the alveolar space, with a concomitant reduction of inflammatory cytokines. This study has delineated the major subsets of lung macrophages driving inflammatory and anti-inflammatory cytokine production within the alveolar space during SARS-CoV-2 infection.

**One sentence summary:** Multi-omic analyses of hyperacute SARS-CoV-2 infection in rhesus macaques identified two population of infiltrating macrophages, as the primary orchestrators of inflammation in the lower airway that can be successfully treated with baricitinib

## INTRODUCTION

The COVID-19 pandemic began with a series of reports of localized outbreaks of pneumonia caused by a novel coronavirus, SARS-CoV-2, in Wuhan, China in December 2019^1,2^. As of late 2021, there have been over 200,000,000 documented infections, and over 4,500,000 fatalities attributed to sequelae of COVID-19. The rapid development and availability of effective vaccines^3–5^ against SARS-CoV-2 infection has provided much needed optimism that infection rates will decline and that the containment of the virus at the population level is possible. Despite these landmark achievements, continued research efforts are essential to safeguard against potential breakthrough variants, to develop therapies for those afflicted while the vaccine rollout continues, and to prevent or minimize the impact of future viral outbreaks. In this light, basic research into the innate and adaptive immune responses to SARS-CoV-2 continues to be critical for informing vaccine and therapeutic approaches directed at ending the COVID-19 pandemic or at decreasing mortality.

Since the emergence of the COVID-19 pandemic, research into the virology, immune responses and pathogenesis of SARS-CoV-2 infection has amassed at an unprecedented rate, and numerous hypotheses have arisen to explain the underlying mechanisms of severe COVID-19. Of these, the concepts that have accumulated the most supporting evidence are: (i) evasion or impairment of early Type I interferon (IFN) responses^6^, (ii) vascular complications arising from hypercoagulability syndromes^7^, and (iii) perturbations of the granulocyte and myeloid compartments in the lower airway and blood manifesting in inflammatory cytokine production^8,9^. Immunologically, severe disease in COVID-19 patients has been associated with a widespread increase in levels of inflammatory mediators (e.g. CXCL10, IL-6, and TNFα) in plasma and bronchoalveolar lavage (BAL) fluid in what is commonly referred to as a “cytokine storm”^10^, and an expansion of macrophages, neutrophils and lymphocytes in the lower airway^8^. Despite this impressive accruement of data, the precise early immunological events and immune cell infiltration that drive inflammation in the lower airway remain uncharacterized.

Non-human primate (NHP) models of SARS-CoV-2 infection (primarily macaque species and African green monkeys (AGMs)) have proven to be critical tools, primarily due to the ability to examine early events after infection longitudinally and in tissues not available in most human studies^11^. NHPs support high levels of viral replication in the upper and lower airway^12–14^, share tissue distribution of ACE2 and TMPRSS2 with humans^15^, and have been invaluable pre-clinical models of vaccines^16–18^ and therapeutics^19,20^. Additionally, mild to moderate COVID-19 has been shown to be recapitulated in SARS-CoV-2-infected NHPs^11^ that typically resolve by 10-15 days post infection (dpi)^11,20,21^. Mechanistic studies of SARS-CoV-2 infection in NHPs have utilized a variety of high-throughput techniques and have reported (i) Type I IFN responses are robustly induced in blood and the lower airway very early after infection^20,22^, (ii) elevated pro-inflammatory cytokines consistent with the “cytokine storm” seen in humans are detectable in plasma and BAL^23^, (iii) vascular pathology and gene expression consistent with hypercoagulability are evident in the lower airways^22^, and (iv) increased production of inflammatory cytokines by myeloid origin cells^20,24^.

In the current study, we used SARS-CoV-2 infected rhesus macaque (RM) to investigate the early inflammatory events occurring in the blood and lower airway using high dimensional flow cytometry, multi-analyte cytokine detection, and bulk and single-cell RNA-Seq (sc-RNA-Seq). To dissect the role of discrete immune subsets within the myeloid fraction in SARS-CoV-2-driven inflammation, we used two different strategies, employing scRNA-Seq and bulk-RNA-Seq reference datasets to classify the macrophage/monocyte population. With this approach, we identified the main subsets of pro-inflammatory macrophages that expand after SARS-CoV2 infection and are the predominant source of inflammatory cytokines in the airway. We also observed an early induction of plasmacytoid dendritic cells (pDCs) in blood and the lower airway that coincided with the peak of the IFN signaling. Finally, we described that treatment of SARS-CoV-2 infected RMs with baricitinib, a JAK1/2 inhibitor recently demonstrated to reduce hospitalization time and mortality for severe COVID-19 patients^25^, suppressed airway inflammation by abrogating the infiltration of pro-inflammatory macrophages to the alveolar space. Collectively, this study defines the early kinetics of pDC recruitment and Type I IFN responses, and identifies discrete subsets of infiltrating macrophages as the predominant source of pro-inflammatory cytokine production in SARS-CoV-2 infection.

## RESULTS

### Study Overview

An overview of the study design is shown in **Fig. 1a**. Eight RMs (mean age 14 years old; range 11-17 years old) were inoculated intranasally and intratracheally with 1.1 × 10^6^ plaque-forming units (PFU) of SARS-CoV-2 (2019-nCoV/USA-WA1/202). These animals were previously reported in a study evaluating the therapeutic efficacy of baricitinib in SARS-CoV-2 infection^20^. At 2dpi, four of the eight animals started receiving baricitinib^20^. For this study, pre-infection baseline and hyperacute time points (1-2dpi) include n = 8 RMs, all untreated, and the remaining longitudinal time-points assessed to determine the pathogenesis of SARS-CoV-2 infection are comprised of n = 4 of the RMs that remained untreated. Inoculation with SARS-CoV-2 led to reproducibly high viral titers detectable in the upper and lower airways by genomic and sub-genomic qPCR assays (**Fig. 1b**). The peak of viremia in the nasal passage, throat and BAL was at 2-4dpi (**Fig. 1b**).

**Figure 1.**
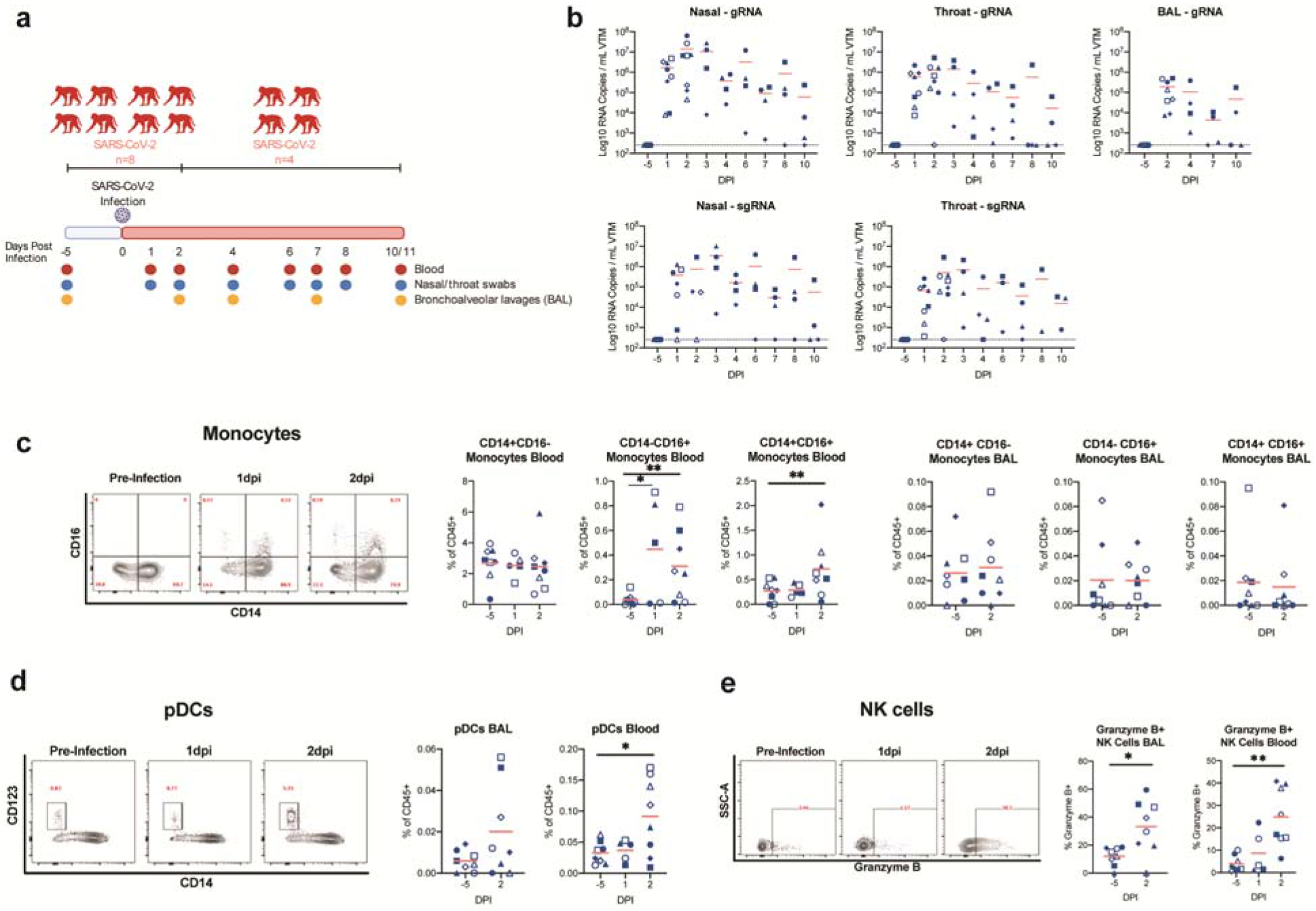
Early expansion of inflammatory cells in the blood following infection with SARS-CoV-2. (**a**) Study design; 8 RMs were infected intranasally and intratracheally with SARS-CoV-2 and tracked longitudinally. Baricitinib was administered daily to 4 RMs starting at 2dpi and the remaining 4 RMs were untreated. (**b**) After SARS-CoV-2 inoculation, nasal, throat, and bronchoalveolar lavages (BAL) were collected and viral loads were quantified by qRT-PCR for total gRNA and sgRNA. (**c**) Longitudinal levels of monocytes within BAL and blood depicted as a % of CD45+ cells. (**d**) Longitudinal levels of plasmacytoid dendritic cells (pDCs) within BAL and blood depicted as a % of CD45+ cells. (**e**) Longitudinal levels of NK cells expressing Granzyme B in BAL and blood. Open symbols represent RMs that received baricitinib treatment starting 2dpi and filled symbols represent untreated RMs. The red bars represent the mean. Statistical analysis was performed using one-tailed Wilcoxon signed-rank test comparing each timepoint to −5dpi. * p-value < 0.05, ** p-value < 0.01.

### SARS-CoV-2 induces a robust, but transient, expansion of pDCs during hyperacute infection

To characterize the innate immune response following SARS-CoV-2 infection, we analyzed changes in innate populations using multi-parametric flow cytometry in blood and BAL samples in the first 2dpi, or “hyperacute” phase of infection (**Fig. 1c-e, Fig. S1**), and over the full course of infection (**Fig. S2**). In blood, we did not observe a significant increase in the proportion of classical monocytes (CD14+CD16-) at 2dpi (**Fig. 1c**) nor at extended time-points (**Fig. S2a)**. Similar to reports in humans^9^, we observed a rapid, but transient, increase in blood CD14-CD16+ and CD14+CD16+ monocytes (**Fig. 1c**, **Fig. S2c)**. Using these conventional markers for blood monocyte subsets, we did not observe any significant changes in CD14-CD16+, CD14+CD16+, nor CD14-CD16+ within the BAL (**Fig. 1c, Fig. S2c)**.

We observed a significantly elevated level of pDCs in blood at 2dpi and similarly, a trend of elevated pDCs in BAL samples (**Fig. 1d, Fig. S2c**). This expansion was transient, as pDC numbers returned to baseline by 4dpi. While the overall frequencies of natural killer cells (NK) were not changed in blood or BAL (**Fig. S2b)**, the fraction of Granzyme B+ NK cells increased significantly at 2dpi in blood, from 4% to 25% **(Fig. 1e)** and remained elevated throughout the course of infection (**Fig. S2**). Similarly, increases in NK cell activation were also observed in the BAL, rising from 12% to 33% at 2dpi **(Fig. 1e)**, and persisting at this level until the study termination at 10/11dpi (**Fig. S2b**). Collectively, these data indicate that during the hyperacute phase of SARS-CoV-2 infection, there is a significant mobilization of innate immune cells capable of initiating and orchestrating effector responses of the Type I IFN system.

### SARS-CoV-2 infection drives robust, but transient, upregulation of IFN responses in blood and lower airway

To understand the extent of immunological perturbations induced by SARS-CoV-2 infection, we performed extensive gene expression profiling of PBMC and BAL samples. During the hyperacute phase, the BAL had widespread induction of pathways associated with innate immunity and inflammation (**Fig. 2a**). Notably, we observed a rapid and robust induction of interferon-stimulated genes (ISGs) in the PBMC and BAL compartments starting at 1 or 2dpi (**Fig. 2b, Fig. S3a**). The ISG response, although widespread, had largely returned to baseline by 10/11dpi (**Fig 2b, Fig. S3a**). We also detected a trend of elevated IFNα protein in 4/6 and 5/8 animals in BAL and plasma, respectively (**Fig. 2c,d)** and a significant increase in RNA-Seq read counts mapping to IFNA genes at 2dpi in BAL (**Fig. 2e)**, which coincided with the expansion of pDCs in the airway and blood (**Fig. 1d**). A significant enrichment of genes representing NK cell cytotoxicity (**Fig. 2a**) was observed at 2dpi in BAL, consistent with our observation of elevated Granzyme B+ NK cells by flow cytometry (**Fig. 1e**). Taken together, these data demonstrate the presence of primary cells able to produce Type I IFNs (i.e., pDCs), coincident with detectable IFNA transcripts and protein, and with downstream IFN-induced effector functions (ISGs, NK cell activation) following SAR-CoV-2 infection, and that these responses were transient, having largely subsided by 10/11dpi.

**Figure 2.**
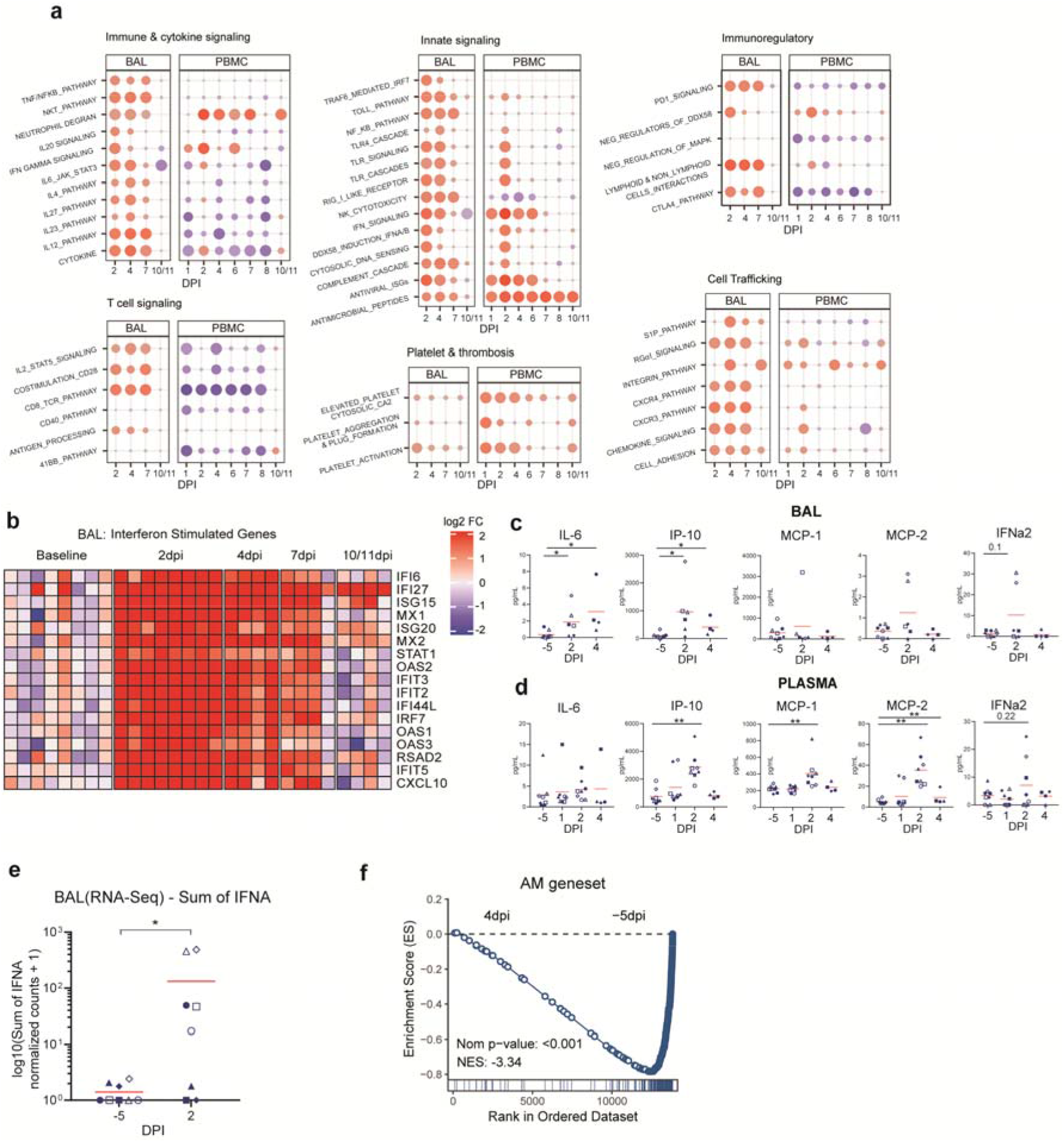
Early pro-inflammatory and ISG response observed in airways and peripheral blood by bulk transcriptomics. **(a)** Dot plots showing normalized enrichment scores and nominal p-values for gene sets. Enrichment is indicated by dot color (red: positively enriched vs −5dpi; blue: negatively enriched), dot size indicates significance. **(b)** Heatmap of longitudinal responses for the ISG gene set. The color scale indicates log2 expression relative to the median of the −5dpi samples. **(c)** Cytokines evaluation (Mesoscale) in BALF and **(d)** Plasma; only significant cytokines are shown. **(e)** Sum of normalized expression of all IFNA genes in BAL. **(f)** GSEA enrichment plot showing negative enrichment for AM gene signature (derived from SingleR) when comparing bulk BAL RNA-Seq samples from 4dpi to −5dpi. The red bar represents the mean. Statistical analysis was performed using one-tailed Wilcoxon signed-rank test comparing each timepoint to −5dpi. * p-value < 0.05, ** p-value < 0.01.

### SARS-CoV-2 infection drives a shift in airway macrophage populations

We observed that SARS-CoV-2 infection induced significant enrichment of several inflammatory cytokine signaling pathways, namely IFNA, IL4, IL6, IL10, IL12, IL23 and TNF, and the chemokine pathways CXCR4 and CXCR3, in both PBMCs and BAL of RMs, with higher magnitude in the BAL (**Fig. 2a, Supplementary Files 1&2**). For many of these pathways, we were able to quantify significant increases in the upstream regulator at either the protein, or mRNA level, or both: IL6 protein levels were significantly increased in the BAL fluid (BALF) (**Fig. 2c**), as were RNA transcripts in BAL (**Fig. S3b)**. Similarly, the induction of CXCR3 pathways signaling was consistent with detection of increased IP10/CXCL10 protein in BALF and RNA at 2dpi in BAL **(Fig. 2c, Fig. S3b)**. The appearance of inflammatory pathways in the blood and airway have been reported in a multitude of human studies (reviewed in^26^). However, we noted that SARS-CoV-2 infection also drove early expression of several immunoregulatory/immunosuppressive pathways in the BAL, namely: PD1 and CTLA4 signaling, and negative regulators of MAP kinase and DDX58/RIG-I signaling (**Fig. 2a)**. Previously, we reported that the myeloid fraction in BAL was primarily responsible for the production of pro-inflammatory mediators, however the specific immunophenotypes were not defined. To further investigate the presence of different macrophage subsets within the lower airway after SARS-CoV-2 infection, we performed GSEA on bulk BAL data using AM gene signature (obtained from SingleR^27^) specific for RM pulmonary macrophages. We observed that genes specific for alveolar macrophages (AMs) were significantly enriched at baseline (−5dpi) relative to 4dpi, indicating a downregulation of this gene set after SARS-CoV-2 infection (**Fig. 2f**). Collectively, these bulk RNA-Seq data indicate a rapid and significant shift in the balance of macrophage populations in the lower airway following SARS-CoV-2 infection.

### SARS-CoV-2 infection induces an influx of two subsets of infiltrating macrophages into the alveolar space

In our prior work in RMs, we demonstrated that cells of myeloid origin were the predominant subset responsible for production of inflammatory cytokines in the lower airway following SARS-CoV-2 infection^20^. While our prior sc-RNA-seq analyses determined the majority of cells in the BAL after infection to be of monocyte/macrophage origin, with relatively few neutrophils or granulocytes, the precise immunophenotypes of the myeloid cells driving inflammation in the lower airway have not been precisely delineated.

Cell classification based on cell-surface marker genes is typically problematic in sc-RNA-seq data due to gene dropouts inherent to the technology. Accurate classification is further complicated in the rhesus model system, in which genomic references have incomplete annotation, and markers from other model species may not phenocopy. Several significant advances have been made recently elucidating the resident tissue macrophage subsets in the lung and their function during viral infection and inflammation^28–31^. However, analysis of sc-RNA-Seq data from RM lung suspensions and BAL during steady state condition indicated that several key markers used to differentiate macrophages in the murine lung (e.g. Lyve1) were not expressed at levels sufficient to distinguish populations in the rhesus pulmonary myeloid populations (**Fig. S4**). Therefore, we used two overlapping alternative strategies to accurately classify tissue macrophages and monocyte-derived/infiltrating macrophages in the RM airway after SARS-Cov-2 infection in our sc-RNA-Seq data. The first strategy was based on using existing lung scRNA-Seq data from uninfected RMs as a reference to map and annotate the BAL cells. We processed lung 10X data from three uninfected RMs (NCBI GEO: GSE149758)^32^ through Seurat pipeline ^33^ and reproduced the four reported macrophage/monocyte subsets: CD163+MRC1+, resembling alveolar macrophages; CD163+MRC1+TREM2+ macrophages, similar to infiltrating monocytes; CD163+MRC1-, similar to interstitial macrophages; and CD16+ non-classical monocytes (**Fig S4 a-d**). We used Seurat to map BAL macrophages/monocytes from SARS-CoV2 infected RMs and transfer annotations from the lung reference. The second strategy involved using bulk RNA-Seq on sorted AM and IM from the lungs of three uninfected RMs, according to the phenotype defined by Cai et al^34^, based on expression of CD206/MRC1 and CD163, to annotate cells using SingleR^27^ **Fig. S4e**). 2069 genes were found to be differentially expressed between IMs and AMs (FDR< 0.05, fold-change>2) (**Fig. S4f**). Of note, CX3CR1 was highly expressed in the IMs, consistent with both murine and human definitions of this subset (**Fig. S4g**). APOBEC3A, an RNA-editing cytidine deaminase, was also highly expressed in IMs along with PTGS2, a pro-inflammatory COX-2 cyclooxygenase enzyme, TIMP1, which enables migration of cells via the breakdown of connective tissue, VCAN, an immunosuppressive regulator, and PDE4B, which regulates expression of TNFα (**Fig. S4g**). We annotated the lung macrophage/monocyte subsets using the bulk sorted AM and IM datasets and found that almost all of CD163+MRC1+ cluster and some CD163+MRC1+TREM2+ cells were annotated as AM and the remaining as IM (**Fig S4h**). Thus, benchmarking our lung sc-RNA-seq based reference against rudimentary bulk transcriptomic signatures demonstrated their accuracy in resolving the AM phenotype from non-AM in steady state conditions.

We next analyzed changes in the myeloid populations within the BAL of RMs after SARS-CoV-2 infection. Using our lung/sc-RNA-seq reference, we found that most of the BAL macrophages/monocytes belonged to the AM-like CD163+MRC1+ macrophage subset at −5dpi along with some cells from the CD163+MRC1+TREM2+ macrophage subset (**Fig. 3ab**). At 4dpi, there was an influx of both CD163+MRC1+TREM2+ macrophages and the IM-like CD163+MRC1-macrophages with few cells annotated as CD16+ non-classical monocytes. The expression of gene markers such as MARCO, FABP4 and CHIT1 further supported the cell subset annotations (**Fig 3c**). We also observed a similar increase in APOBEC3A and decreases in MARCO and CHIT1 expression in bulk BAL samples (**Fig 3d**). The percentage of CD163+MRC1+ macrophages reduced overall from 70% to 29% of all macrophages/monocytes in BAL (**Fig 3e**). Similarly, we saw an overall increase in the percentage of CD163+MRC1+TREM2+ macrophages from 5% to 21% and the IM-like CD163+MRC1-macrophages from 0.1% to 4%. Thus, COVID-19 infection resulted in an influx of monocyte-derived and IM-like macrophages in BAL at 4dpi. We also found that these infiltrating macrophages expressed higher levels of several pro-inflammatory cytokines and chemokines compared to the CD163+MRC1+ AM-like macrophages (**Fig 3f-h, Supplementary File 4**).

**Figure 3.**
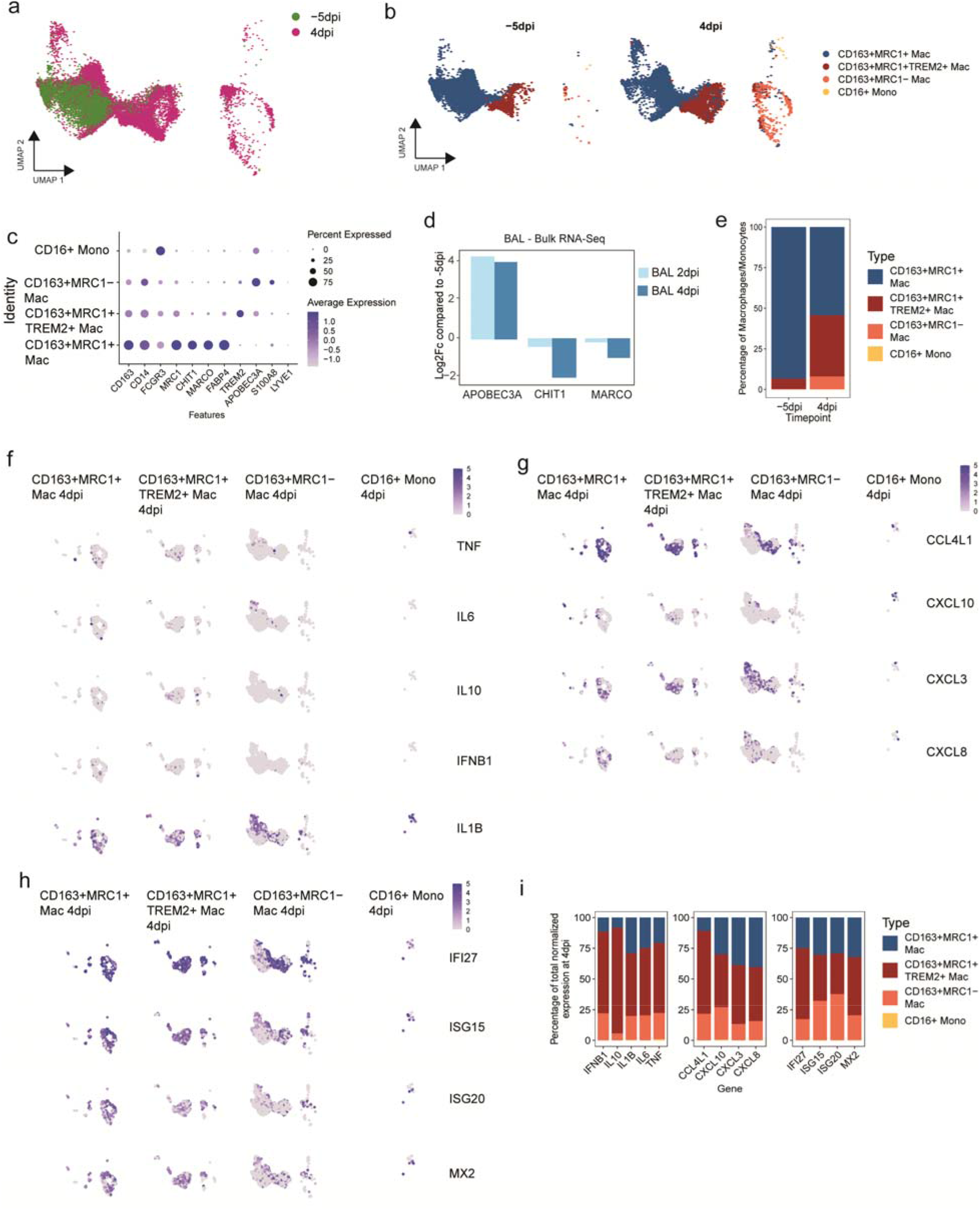
Influx of pro-inflammatory macrophages in BAL. **(a)** Projection of single-cell macrophages/monocytes from −5dpi (green) and 4dpi (magenta) 10X BAL samples obtained from three SARS-CoV2 infected rhesus macaques onto the reference UMAP of lung macrophage/monocytes from uninfected rhesus macaques (NCBI GEO : GSE149758). **(b)** UMAP projections showing the predicted cell type annotations from the uninfected lung reference split by time of sample collection. **(c)** DotPlots showing the expression of marker genes for the different macrophage/monocyte subsets in SARS-CoV2 infected BAL samples **(d)** Log2 fold-changes compared to −5dpi for APOBEC3A, CHIT1 and MARCO in bulk BAL RNA-Seq data. **(e)** Percentage of a given subset out of all macrophage/monocyte subsets at −5dpi and 4dpi from all three animals **(f,g,h)** FeaturePlots showing the expression of selected pro-inflammatory cytokines (f), chemokines (g) and ISG (h) in different macrophage/monocyte subsets at 4 dpi. **(i)** Contribution of each macrophage/monocyte subset towards the production of the pro-inflammatory genes and ISG. The percentage contribution was calculated by dividing the sum of normalized expression of a given gene in a macrophage/monocyte subset by the sum of the normalized expression of the gene in all macrophage/monocyte subsets.

To further validate our cell classification and support the observation that it is the infiltrating cells that increase in numbers and predominantly producing inflammatory mediators, we used the second strategy of using gene expression of bulk sorted AM and IM cells to classify the BAL macrophages/monocytes. Using this definition, we confirmed that there is an increase in the percentage of non-AM population with a corresponding decrease in the AM population (**Fig S5b, d**). The non-AM population was also found to show higher expression of pro-inflammatory cytokines (**Fig S5e, Supplementary File 4**).

Differential expression analysis of 4dpi and −5dpi BAL macrophages/monocytes showed that CHIT1, MARCO and MRC1 were among the top-ranking genes exhibiting downregulation in the BAL, while genes such as ADAMDEC1^35^ and S100A8^36^ that are associated with monocyte-derived macrophages were among the most upregulated (**Supplementary File 4**). These data demonstrate that our observation of an influx of infiltrating macrophages into the BAL at 4dpi was consistent across multiple definitions of this phenotype.

### Infiltrating macrophages produce the majority of lower airway inflammatory cytokines during acute SARS-CoV-2 infection

Given our observation of the dynamics of pulmonary macrophages within the alveolar space during early SARS-CoV-2 infection, we characterized the transcriptional changes in each macrophage/monocyte population. Several chemokines (CCL4L1, CCL3, CXCL3, CCL2), multiple ISGs, NFKB1A, S100A8, and GZMB were among the most upregulated genes at 4dpi in BAL populations (**Supplementary File 4**). Elevated expression of multiple inflammatory genes, including IL6, TNF, IL10, IFNB1 and IL1B, were observed in the CD163+MRC1+TREM2 Mac and CD163+MRC1-subsets (**Fig. 3f, Fig. S6**) after infection. The infiltrating macrophages were also observed to upregulate multiple chemokines, including those specific for recruiting neutrophils (CXCL3, CXCL8), macrophages (CCL2, CCL3, CCL5, CCL4L1), and activated T cells (CXCL10) as well as multiple ISGs (**Fig. 3g-h, Fig. S6)**. When we examined CD163+ MRC1+ macrophages, many of the same inflammatory cytokines and gene sets seen in the infiltrating macrophages were elevated at 4dpi, albeit at much lower magnitude (**Fig. 3f-h, Fig. S6**). Having observed a significantly higher average expression of inflammatory cytokines in infiltrating macrophages compared to CD163+MRC1+ macrophages, we compared the fractions of sequencing reads detected from each of the subsets to assess the overall contribution to inflammatory cytokine production (**Fig. 3i**). At 4dpi, we observed that the CD163+MRC1+TREM2+ macrophages accounted for 55% of IL6, 57% of TNF, 86% of IL10 and 66% of IFNB1 expression while the CD163+MRC1-macrophages accounted for 20% of IL6, 21% of TNF, 6% of IL10 and 19% of IFNB1 expression. We also found that CD163+MRC1-macrophages expressed higher levels of several of these genes (**Fig. S6**). Thus, infiltrating macrophages are responsible for the majority of lower airway inflammatory cytokine production during acute SARS-CoV-2 infection.

### Baricitinib treatment prevents the influx of inflammatory IM into the lower airway

Baricitinib is a JAK1/2 inhibitor approved for the treatment of active rheumatoid arthritis that was recently granted emergency use authorization for the treatment of hospitalized COVID-19 patients, and reported to reduce mortality when administered as monotherapy^37^ or in combination with remdesivir^25^. In our earlier study, we found that baricitinib was able to suppress the expression of pro-inflammatory cytokines in BAL of RMs infected with SARS-CoV-2^20^. Here, we extended this study to further characterize the impact of baricitinib on the myeloid populations in the airway from five RMs before infection (−5) and at 4dpi, with three RMs that remained untreated and two that received baricitinib). We found that two days of baricitinib administration virtually abrogated the influx of infiltrating macrophages into the alveolar space at 4dpi (**Fig. 4a-c**). This observation was consistent using classifications of macrophages either based on mapping to 10X lung reference or using bulk sorted AM/IM cells (**Fig. 4a-c, Fig. S5a-d**). In addition to preventing the influx of infiltrating macrophages, baricitinib treatment also resulted in significantly lower expression of inflammatory cytokines and chemokines, but the ISG expression remained comparable to untreated animals (**Fig. 4d-f, Fig. S5e**). In summary, these data further elucidate the mechanism of action by which baricitinib treatment abrogates airway inflammation in SARS-CoV-2 infection^20^, by demonstrating its ability to block infiltration of discrete pro-inflammatory macrophage populations into the alveolar compartment.

**Figure 4.**
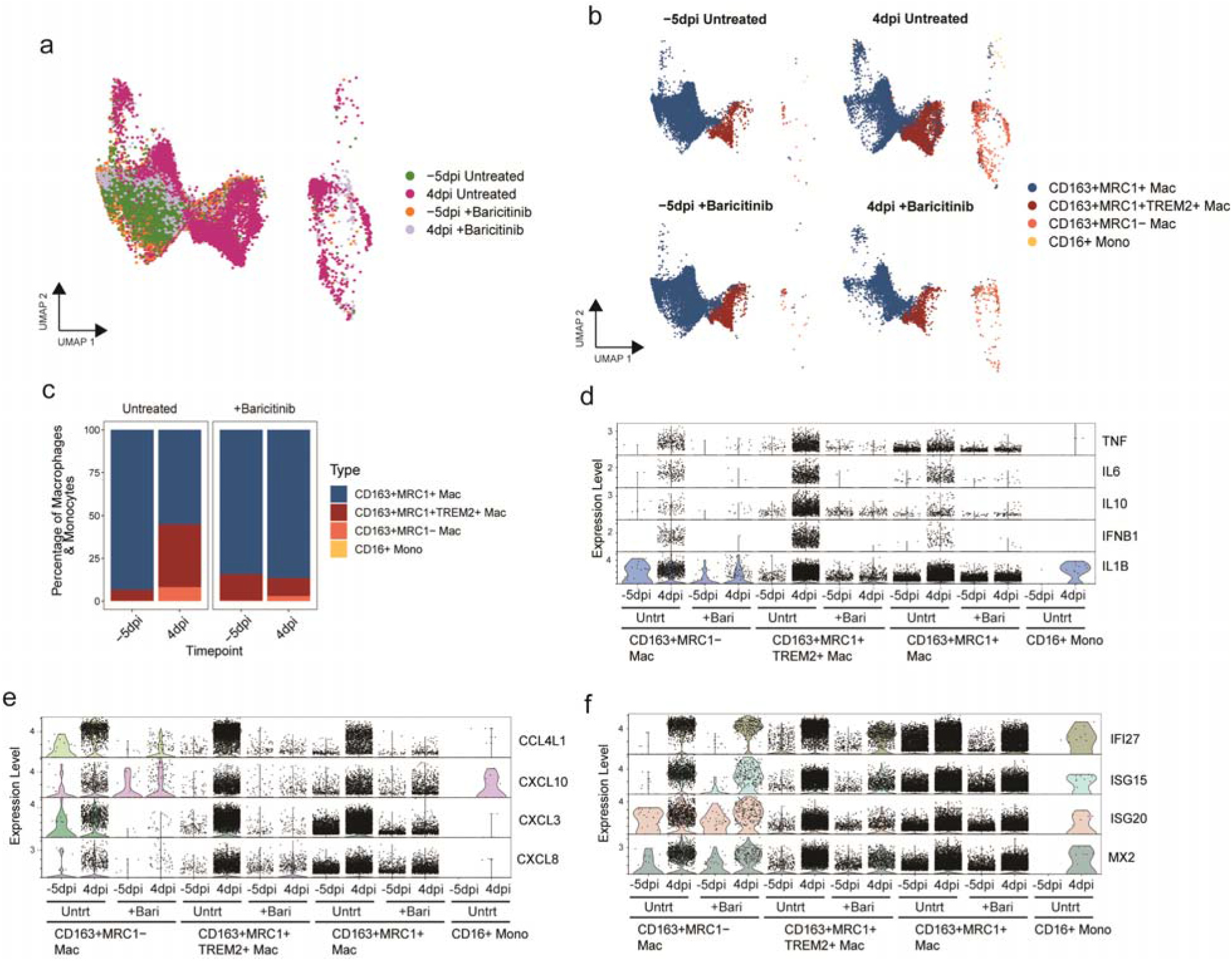
Baricitinib reduced the influx of pro-inflammatory macrophages in addition to the pro-inflammatory gene expression profile. **(a)** Projection of macrophages/monocytes from −5dpi and 4dpi 10X BAL samples from three untreated and two baricitinib treated rhesus macaques on the reference UMAP of uninfected lung macrophages/monocytes (NCBI GEO: GSE149758) **(b)** UMAP split by treatment and timepoint showing predicted cells annotations based on mapping to the reference lung macrophages/monocytes **(c)** Percentage of a given macrophage/monocyte subset of all the macrophages/monocytes in the BAL samples. **(d)** Violin plots showing expression of pro-inflammatory cytokines, chemokines and ISG in the different macrophage/monocyte subsets in BAL 10X samples from baricitinib treated and untreated samples.

## DISCUSSION

The mechanisms by which SARS-CoV-2 infection establishes severe disease remain largely unknown, but remain a key priority for reducing the toll of the COVID-19 pandemic. As the appearance of symptoms range from 2-14 days after SARS-CoV-2 infection, characterization of the early immunological events using clinical samples is challenging. Here, we utilized the RM model of SARS-CoV-2 infection and an integrated systems analysis to dissect the immune response during hyper-acute infection. Our findings were: (i) SARS-CoV-2 infection initiated a robust Type I IFN response in the blood and lower airway apparent at 1-2dpi; (ii) SARS-CoV-2 induced a rapid influx of two infiltrating macrophage populations, into the bronchoalveolar space, which produced the majority of inflammatory cytokine production; and (iii) the mechanism of action of baricitinib, a drug recently authorized for emergency use in the treatment of severe COVID-19, is to abrogate infiltration of these inflammatory cells into the airway. Our data present, to date, the most comprehensive analysis of the immunopathological events occurring during hyperacute SARS-CoV-2 infection.

Using our reference datasets of RM lung macrophages, we identified two myeloid cell subsets, both clearly distinct from alveolar macrophages, infiltrating the airway after SARS-CoV-2 infection, that were the main producers of lower airway inflammatory cytokines and chemokines. One population, defined as CD163+MRC+TREM2+ cells, were highly similar to murine definitions of infiltrating CCR2+ monocytes. The second, CD163+MRC-, largely resembled interstitial macrophages. Our data are consistent with a recent observation of a rapid (3dpi) increase of IMs in the BAL of RMs using flow cytometry^21^. Similarly, an accumulation of non-AMs (defined as CD16+CD206-HLA-DR+/CD11b+), and reciprocal reduction of AMs, has been observed in the BAL and lungs of infected RMs and AGMs^23^. Lastly, our data are consistent with our recent findings in the murine model, in which SARS-CoV-2 elicited recruitment of circulating monocytes to the lung parenchyma, but was significantly abrogated in CCR2-deficient mice^38^. The CD163+MRC1+ Mac/AM-like subset also contributed to the inflammatory milieu, producing IL6, TNF, and IL10, albeit in significantly lower quantities.

It is important to note that our observations were during hyperacute infection, and that our animals did not develop severe disease, so although our data indicate that these infiltrating populations orchestrate early inflammation and may contribute to airway pathogenesis, we cannot formally make this link. However, this model is consistent with recent data by Ren et al^39^., who observed a significant loss of MARCO expression in BAL-resident myeloid populations of patients with severe COVID-19 relative to those with moderate disease, similar to our observations, in which the appearance of infiltrating macrophages diluted the population of MARCO+ macrophages^40^. Those observations, taken together with our data, suggest that the inflammatory macrophage phenotype we identify here may be preferentially retained in the lower airway of patients with severe COVID-19. Additionally, we demonstrated that *in vivo* treatment with the JAK1/2 inhibitor baricitinib, which has demonstrated efficacy in reducing severe disease, was able to virtually abrogate the recruitment of these inflammatory macrophages into the airway, providing an additional mechanistic link with the development of COVID-19-related pathogenesis.

In addition to inflammatory cytokines, we observed that the infiltrating macrophage subsets produced high levels of IL10, and were enriched in IL10 signaling pathways. Lung IM’s are considered to be a “professional IL10-producing cell” producing IL10 at both a steady state and in response to innate stimuli (LPS, unmethylated CpGs)^41^. The majority of data to date has demonstrated an immunoregulatory, protective role for IMs in murine models of asthma, lung fibrosis, and allergen induced inflammation^29^. However, while the pro-inflammatory potential of IMs has been relatively understudied, they have been demonstrated to be efficient at producing IL6 and TNF in response to TLR ligands^42^. Given our observations of high IL10 production in infiltrating macrophages, we cannot exclude a potential immunoregulatory role for this subset, and indeed, it presents an interesting hypothesis in which the balance of infiltrating IM vs TREM2+ macrophages into the bronchoalveolar space determines the pathogenic outcome of SARS-CoV-2 infection. Lastly, recent publications have reported that lung IMs may be comprised of two, or even three, functionally distinct populations, defined by an axis of expression of Lyve1, MHC, CD169, and CD11c^28–31^. We did not observe separate clustering amongst BAL IMs, nor differential expression amongst these markers, and further work is needed to understand the congruency of macaque macrophage subsets with those identified in the murine model.

We observed a very rapid and robust induction of the Type I IFN pathway at 1-2dpi, characterized by elevated pDCs in the airway and blood, IFNA and IFNB transcripts and protein, upregulated ISGs, and increased granzyme B in NK cells. The Type I IFN response in SARS-CoV-2 infection has been intensely studied: in vitro infection of airway epithelial cells have consistently resulted in a muted ISG response^43^, and patients developing severe COVID-19 have been reported to have higher incidence of mutations in IFN response genes, or elevated levels of autoantibodies against IFN-response genes (reviewed in^6,44–51^). Our data, in which the IFN response peaked at 2dpi and had largely abated by 10/11dpi, provides well defined kinetics of the ISG response, and similar observations have been reported in other NHP studies^21,22^. The rapid and short-lived nature of the IFN response underscores the difficulty in interpreting the IFN response in clinical samples.

Our multiparametric analyses demonstrated an increase of pDCs at 2dpi that coincided with the peak of ISG production, IFNA/B detection, and NK cell activation, thus implicating pDCs as the primary cell orchestrating the IFN response in the lower airway. We had previously observed a reduction in peripheral blood pDCs frequencies and activity in human SARS-CoV-2 infection^52^, and other have reported signatures of pDCs apoptosis that predicted lower IFN-I responses^53^. Taken in the context of these clinical findings, our observation of pDC accumulation in the BAL indicates that they undergo rapid mobilization from the blood to the lower airway, and this suggests they likely drive early protective innate immune responses. However, pDCs may also contribute to pathological inflammation; and future interventional studies targeting the pDC/IFN axis in animal models will be necessary to test these hypotheses.

In our prior study, we demonstrated the ability of baricitinib to block airway pro-inflammatory cytokine production in SARS-CoV-2-infected RMs while preserving Type I IFN responses^20^. Here, we extended these findings to demonstrate that baricitinib blocked the influx of inflammatory macrophages into the bronchoalveolar space. These data adds to our mechanistic understanding of the action of baricitinib, and provide a potential explanation for the disparity of baricitinib’s impact on IFN vs IL6/TNF signaling when considering the timing of the drug administration. We administered baricitinib at 2dpi, after the peak influx of pDCs, but before the likely appearance of the APOBEC3A+ inflammatory macrophages at 3-4dpi. The ongoing ISG response, and suppressed TNF/IL6 response, suggest that the primary mechanism by which baricitinib protects the airway is by blocking recruitment of inflammatory cells to the bronchoalveolar space. In regards to guiding future clinical application of baricitinib, our data suggest that timing is critical, and would favor earlier drug administration.

Our study had some limitations; first, while the RM/SARS-CoV-2 model has rapidly been adopted by several groups for pre-clinical testing of anti-COVID drugs and vaccines, no group has demonstrated overt, reproducible symptomatic disease^11^. Thus, linking early immunological events to the development of severe COVID-19 requires validation in human studies, such as the observations of reduced MARCO expression in the airway myeloid populations of severe COVID-19 patients noted above^40^. Another drawback was the relatively low power of our study. While our observations at 0, 1, and 2dpi were n = 8, we were limited to n = 4 for day 4-10 observations. However, it should be noted that there was no lack of statistical power for our key observations.

While the global vaccine rollout has made great strides to reduce the transmission and severity of SARS-CoV-2 infection, millions of people remain vulnerable. Understanding the early events of SARS-CoV-2 infection, and the mechanisms by which clinically approved drugs afford protection, remains a global priority. In this study, we have identified a novel population of inflammatory myeloid cells that are responsible for the preponderance of airway inflammation in acute SARS-CoV-2 infection. We also demonstrated that treatment with the emergency authorized JAK1/2 inhibitor baricitinib in combination with remdesivir, blocked infiltration of these inflammatory cells into the alveolar space. These data identify both a key druggable target (airway infiltrating macrophages), and an efficacious mechanism by which to lower airway inflammation, and should prove useful for identifying additional drugs to reduce the incidence and mortality of severe COVID-19 disease.

## MATERIALS AND METHODS

### Animal and SARS-CoV-2 infections

The animal care facilities at YNPRC are accredited by the Association for Assessment and Accreditation of Laboratory Animal Care (AAALAC) as well as the U.S. Department of Agriculture (USDA). Emory University’s Institutional Animal Care and Use Committee (IACUC) reviewed and approved all animal experiments under permit PROTO202000035. All procedures were performed as per the institutional regulations and guidelines set forth by the NIH’s Guide for the Care and Use of Laboratory Animals (8^th^ edition) and were conducted under anesthesia and appropriate follow-up pain management to minimize animal suffering. Eight (4 female, 4 males, aged >11 yrs) specific-pathogen-free Indian-origin rhesus macaques were infected via intranasal and intratracheal routes with 1.1 × 10^6^ plaque forming units (PFU) SARS-CoV-2 (see SI for details), as previously described^20^ and were maintained in the ABSL3 at YNPRC. The processing of nasopharyngeal swabs, BAL and mononuclear cells was performed as described previously^20^.

### Determination of viral load RNA

The SARS-CoV-2 genomic and sub-genomic RNA was quantified in nasopharyngeal swabs, throat swabs, and BAL as previously described^18,20^(see SI for details).

### Immunophenotyping and flow cytometric purification of RM pulmonary macrophages

23-parameter flow cytometric analysis was perform on fresh PBMCs and BAL mononuclear cells from SARS-CoV-2 infected RMs as described previously^20^. For purification of CD163^+^CD206^+^ (AM), CD163^+^CD206^−^ (IM) cells, cryopreserved single-cell lung suspensions from RMs were stained with BD CD163(GHI/61), CD206(19.2)(BD Biosciences), and purified using BD FACSAria in the Regional Biocontainment Laboratory at the University of Pittsburgh.

### Bulk and sc-RNA-Seq Library and sequencing

The data for −5dpi, 2dpi and 4dpi for bulk BAL samples was obtained from our previous study^20^. Here we expanded our study to include 7dpi and 10dpi/11dpi samples for BAL and −5dpi, 1dpi, 2dpi, 4dpi, 6dpi, 7dpi, 8dpi and 10/11dpi for PBMC. Cell suspensions were prepared in BSL3, for bulk RNA-Seq, 250,000 cells (PBMCs) or 100,000 cells (BAL) were lysed directly into 700 ul of QIAzol reagent. Libraries were prepared as described previously^20^ and sequenced on an Illumina NovaSeq6000 at 100SR, yielding 20-25 million reads per sample. For sc-RNA-Seq, approximately 30,000 cells were loaded onto a 10X Chromium Controller in a BSL3 and single cells were partitioned into droplets using Chromium NextGEM Single Cell 50 Library & Gel Bead kits(10X Genomics, Pleasanton, CA). cDNA was amplified and libraries were prepared for sequencing according to manufacturer instructions. Gene expression libraries were sequenced as paired-end 26×91 reads targeting 50,000 reads per cell.

### scRNA-Seq and bulk RNA-Seq analysis

The filtered count matrices for BAL were obtained from^20^. The Seurat library v4.0.4^33^ was used to perform the analysis (see SI for details). The macrophages/monocytes from BAL samples were annotated into subsets using two approaches – (i) mapping to macrophages/monocytes from lung reference using Seurat and (ii) using bulk sorted cells as reference with SingleR^27^. The 10X lung scRNA-seq data from three uninfected macaque was obtained from a published study (NCBI GEO: GSE149758)^32^. The SingleR library was used for cell classification with the BluePrintEncodeData reference. Regularized log counts obtained from DESeq2 for bulk sorted IM and AM were used as reference to annotate the cells from BAL samples using SingleR with default parameters. Bulk RNA-Seq analysis was performed as previously described^20^.

### Mesoscale cytokine analysis

U-PLEX assays (Meso Scale MULTI-ARRAY Technology) were used for plasma and BALF cytokine detection according to manufacturer’s instructions, using 25 microliters as input.

## Data and materials availability

Source data supporting this work are available from the corresponding author upon reasonable request.

## Code availability

The scripts used for analysis are available at https://github.com/BosingerLab/NHP_COVID-19_2.

## Acknowledgments

We kindly thank the Yerkes National Primate Research Center (YNPRC) Director Paul Johnson; Division of Animal Resources, especially Joyce Cohen, Stephanie Ehnert, Stacey Weissman, Sherrie Jean, Jennifer S. Wood, Fawn Connor-Stroud, Rachelle L. Stammen, Racquel A. Sampson-Harley, Denise Bonenberger, John M. Wambua, Dominic M. D’Urso, Sanjeev Gumber, Kalpana Patel, Maureen Thompson, Ernestine Mahar and Kyndal Goss for providing support in animal care, biosafety guidance, technical and computational support. We apologize for key publications omitted due to space limitations.

## Funding

NIH Office of Research Infrastructure Programs (ORIP) P51 OD11132 to YNPRC.

Emory University COVID-19 Molecules and Pathogens to Populations and Pandemics (MP3) Initiative Seed Grant to M.P., A.P., and R.F.S.

YNPRC Coronavirus Pilot Research Project Program grant (to M.Pa. under award P51 OD11132). Fast Grants #2144 to M.Pa

The William I.H. and Lula E. Pitts Foundation Grant to M.Pa.

NIAID award P30 AI050409 to the Center for AIDS Research (CFAR) at Emory University and R01-MH-116695 to R.F.S.

Sequencing data was acquired on an Illumina NovaSeq6000 funded by NIH S10 OD026799 to S. B. This work was additionally funded by the National Institute of Allergy and Infectious Diseases (NIAID, NIH) under awards R37AI141258 and R01AI116379 to M. Paiardini, R01MH116695 to R.F.S, R01AI143411, and R01HL140223 to R.D.L.

## Author contributions

Conceptualization: A.U.,T.H.,M.Pi.,M.Pa, R.F.S & S.B.;

Methodology: T.H., M.Pi., E.V.,S.P.R. M.L.,J.C.,Z.S.,D.C.,E.N.B.,T.Ho.,S.H.,H.A.,J.H.,K.N.,K.P.,A.P.,R.D.,& T.H.V.;

Formal Analysis, A.U.,T.H.,M.Pi.,A.B.,M.L.,Z.S.,E.V.,G.T., & T.H.V;

Investigation: A.U.,T.H.,M.Pi.,M.Pa., & S.B.;

Resources: S.B.B.,R.A.,R.F.S., R.P.S.,M.Pa. & S.B.;

Writing: – Original Draft, A.U., S.B.;

Writing, Review & Editing, T.H.,S.P.R.,M.Pi. M.Pa, R.F.S.

Visualization, A.U.,T.H.,M.Pi., A.B., G.K.T.,.;

Supervision: M.Pa. and S. B.;

Funding Acquisition: T.V.,A.P.,R.F.S.,R.P.S.,M.Pa & S.E.B.

## Competing interests

R.F.S. has served as an unpaid consultant for Eli Lilly whose drugs are being evaluated in the research described in this paper, and owns shares in Eli Lilly. The terms of this arrangement have been reviewed and approved by Emory University in accordance with its conflict of interest policies. All other authors do not have any conflicts to declare.

