## Supplementary Information for "TREM2+ and interstitial macrophages orchestrate airway inflammation in SARS-CoV-2 infection in rhesus macaques"

**List of Supplementary Materials**

**Materials & Methods**

**Supplementary Figure 1.** Flow sorting strategy

**Supplementary Figure 2.** Longitudinal flow cytometric analysis in BAL and blood following SARS-CoV-2 infection.

**Supplementary Figure 3.** Bulk transcriptomic analysis of airways and peripheral blood.

**Supplementary Figure 4** Reference used for annotating macrophage/monocyte subsets.

**Supplementary Figure 5.** Cell annotation using bulk sorted cells as reference

**Supplementary Figure 6**. Expression of canonical marker and pro-inflammatory genes in different macrophage/monocyte subsets.

**Supplementary File 1.** DESeq2 results for bulk BAL samples

**Supplementary File 2.** DESeq2 results for bulk PBMC samples

**Supplementary File 3**. Cell numbers for each macrophage/monocyte subset

**Supplementary File 4.** Differential gene expression analysis for macrophage/monocyte subsets in 10X BAL samples

**Supplementary File 5.** Contribution of macrophage subsets towards inflammatory gene expression

**Material and Methods**

***Viral Stocks***

The viral stocks used for infecting RM were previously described^1^. SARS-CoV-2 (NR-52281: BEI Resources, Manassas, VA; USA-WA/2020, Lot no. 70033175) was passaged on Vero E6 cell line (African Green Monkey Kidney cell line; CRL-1586, ATCC) at a MOI of 0.01. SARS-CoV-2 was propagated and titrated by TCID_50_ method followed by storage of aliquots at -80°C until further use. The infectious dose delivered was determined by back titration of viral stocks via plaque assay. The virus stock was sequenced to confirm the presence of furin cleavage motif. The viral stocks used had less than 6% of genomes with a mutation that may abrogate furin cleavage.

***Determination of viral load RNA***

The swabs were kept in 1mL of Viral Transport Medium (VTM-1L, Labscoop, LLC). The viral RNA was extracted from fresh specimens of nasopharyngeal (NP) swabs, throat swabs, and BAL manually using the QiaAmp Viral RNA mini kit as per the manufacturer’s protocol. For genomic RNA, N2 primer and probe set designed by the CDC for their diagnostic algorithm: CoV2-N2-F: 5’-TTACAAACATTGGCCGCAAA-3’, CoV2-N2-R: 5’-GCGCGACATTCCGAAGAA-3’, and CoV2- N2-Pr: 5’-FAM-ACAATTTGCCCCCAGCGCTTCAG-BHQ-3’^2^ were used for quantitative PCR (qPCR). For sub-genomic RNA, the primer and probe sequences for E gene subgenomic mRNA transcript^3^ were used: SGMRNA-E-F: 5’-CGATCTCTTGTAGATCTGTTCTC-3’, SGMRNA-E-R: 5’-ATATTGCAGCAGTACGCACACA-3’, and SGMRNA-E-Pr: 5’-FAM-ACACTAGCCATCCTTACTGCGCTTCG-3’. The qPCR reactions were performed with the TaqMan Fast Virus 1-step Master Mix using the manufacturer’s cycling conditions, 200nM of each primer, and 125nM of the probe in duplicate. The limit of detection in this assay was 257 copies per mL of VTM/plasma/BAL. The CDC RNase P p30 subunit qPCR, modified for rhesus macaque specific polymorphisms, was used to verify sample quality using the following primer and probe sequences: RM-RPP30-F 5’-AGACTTGGACGTGCGAGCG-3’, RM-RPP30-R 5’-GAGCCGCTGTCTCCACAAGT-3’, and RPP30-Pr 5’-FAM-TTCTGACCTGAAGGCTCTGCGCG-BHQ1-3’. The RNA integrity and sample quality was verified by running a single well from each extraction.

***Tissue processing***

NP swabs were collected under anesthesia by using a clean rayon-tipped swab (ThermoFischer Scientific, BactiSwab NPG, R12300) placed approximately 2-3cm into the nares.  Oropharyneal swabs were collected under anesthesia using polyester tipped swabs (Puritan Standard Polyester Tipped applicator, polystyrene handle, 25-806 2PD, VWR International) to streak the tonsils and back of throat bilaterally (throat/pharyngeal). The swabs were dipped in 1 mL viral transport media (Viral transport Media, VTM-1L, Labscoop, LLC) and vortexed for 30 sec, and the eluate was collected.

To collect BAL, a fiberoptic bronchoscope (Olympus BF-XP190 EVIS EXERA III ULTRA SLM BRNCH and BF-P190 EVIS EXERA 4.1mm) was manipulated into the trachea, directed into the primary bronchus, and secured into a distal subsegmental bronchus upon which 35-50 mL of normal saline (0.9% NaCl) was administered into the bronchus and re-aspirated to obtain a minimum of 20ml of lavage fluid. BAL was filtered through a 70μm cell strainer.

Mononuclear cells were counted for viability using a Countess II Automated Cell Counter (Thermo Fisher) with trypan blue stain and were cryo-preserved in aliquots of up to 2x10^7^ cells in 10% DMSO in heat-inactivated FBS. Whole tissue segments (0.5 cm^3^) were snap frozen dry, or stored in RNAlater (Qiagen), or Nuclisens lysis buffer (Biomerieux) for analyses of compound distribution, RNA-seq, and tissue viral quantification, respectively.

***Immunophenotyping***

The following mAbs were used for the phenotyping of innate immune cells in whole blood (500 μL), as described in^4^, and mononuclear cells (10^6^ cells) derived from BAL: anti-CD20-BB700 (clone 2H7; 2.5 μL; cat. # 745889), anti-Ki-67-BV480 (clone B56; 5 μL; cat. # 566109), anti-CD14-BV605 (clone M5E2; 2.5 μL; cat. # 564054), anti-CD56-BV711 (clone B159; 2.5 μL; cat. # 740781), anti-CD115-BV750 (clone 9-4D2-1E4; 2.5 μL; cat. # 747093), anti-CD3-BUV395 (clone SP34-2; 2.5 μL; cat. # 564117), anti-CD8-BUV496 (clone RPA-T8; 2.5 μL; cat. # 612942), anti-CD45-BUV563 (clone D058-1283; 2.5 μL; cat. # 741414), anti-CCR2-BUV661 (clone LS132.1D9; 2.5 μL; cat. # 750472), anti-CD16-BUV737 (clone 3G8; 2.5 μL; cat. # 564434), anti-CD69-BUV805 (clone FN50; 2.5 μL; cat. # 748763), and Fixable Viability Stain 700 (2 μL; cat. # 564997) all from BD Biosciences; anti-CD38-FITC (clone AT1; 2.5 μL; cat. # 60131FI) from STEMCELL Technologies; anti-CD161-BV421 (clone HP-3G10; 5 μL; cat. # 339914), anti-HLA-DR-BV650 (clone L243; 5 μL; cat. # 307650), anti-CD11c-BV785 (clone 3.9; 5 μL; cat. # 301644), anti-CD11b-PE (clone ICRF44; 2.5 μL; cat. # 301306), and anti-CD123-APC-Fire750 (clone 315; 2.5 μL; cat. # 306042) all from Biolegend; anti-GranzymeB-PE-TexasRed (clone GB11; 2.5 μL; cat. # GRB17) from Thermo Fisher; anti-CD66abce-PE-Vio770 (clone TET2; 1 μL; cat. # 130-119-849) from Miltenyi Biotec; and anti-CD27-PE-Cy5 (clone 1A4CD27; 2.5 μL; cat. # 6607107) and anti-NKG2A-APC (clone Z199; 5 μL; cat. # A60797) from Beckman Coulter. The sorting strategy is show in **Fig. S1**.

***Bulk RNA-Seq library & sequencing***

RNA was isolated using RNeasy Mini or Micro kits (QIAGEN) with on-column DNase digestion. The quality of RNA was determined using an Agilent Bioanalyzer and the cDNA synthesis was carried out using the total RNA with Clontech SMARTSeq v4 Ultra Low Input RNA kit (Takara Bio) as per the manufacturer’s instructions. Dual-indexed bar codes were appended to the amplified cDNA after fragmenting using the NexteraXT DNA Library Preparation kit (Illumina). Agilent 4200 TapeStation was used to validate the libraries by capillary electrophoresis and the libraries were pooled at equimolar concentrations,

***Bulk RNA-Seq analysis***

The STAR index was built by combining genome sequences for Macaca mulatta (Mmul10 Ensembl release 100), SARS-CoV2 (strain MN985325.1 - NCBI) and ERCC sequences as described previously^1^. The ReadsPerGene files were used to generate counts in the htseq format and were imported in DESeq2^5^ using the DESeqDataSetFromHTSeqCount function.

The PBMC and BAL samples were analyzed separately and the design used was: ∼ Group + Subject + Timepoint where Group distinguished between samples that were untreated or treated with baricitinib during the time course. Differentially expressed genes for BAL and PBMC were determined using a threshold of padj < 0.05, fold-change > 2 and filtering out lowly expressed genes where all of the samples at a particular timepoint were required to have detectable expression by normalized reads > 0 for that gene.

In order to obtain references for assigning cell types in single-cell data, bulk RNA-Seq data of interstitial (IM) and alveolar macrophages (AM) from three uninfected rhesus macaques was analyzed using DESeq2. The regularized log expression values were obtained using the rlog function with the parameters blind = FALSE and filtType = “parametric.” The significant genes were filtered based on following criteria: padj < 0.05; fold-change > 2 and normalized mean expression > 5000 for either IM or AM samples.

The input for GSEA was the regularized log expression values obtained from DESeq2. The following gene sets were used for GSEA^6^ analysis: Hallmark and Canonical pathways (MsigDB), NHP ISGs^7^ and Rheumatoid arthritis (KEGG map05323). GSEA was run with default parameters with the permutation type set to gene_set. Volcano plots of differential expression at each timepoint were generated with Enhanced Volcano R library^8^. The regularized log expression values from DESeq2 were used to generate heatmaps using the Complex Heatmap R library^9^.

***scRNA-Seq analysis***

For each BAL sample from SARS-CoV2 infected rhesus macaque, the count matrix was filtered to include only the protein coding genes. Genes encoded on Y chromosome, mitochondrial genes, RPS and RPL genes, B-cell receptor and T-cell receptor genes, and HBB were filtered out. The following parameters were used to filter cells: (i) nFeature_RNA >=200 & <=4000, (ii) % of HBB gene < 10, (iii) % of mitochondrial genes < 20, (iv) % of RPS/RPL genes < 30 and (v) log10(nFeature_RNA) / log10(nCount_RNA) >= 0.8. The number of cells from each sample that passed QC metrics are included in Supplementary File 3. All the BAL samples from each animal at -5dpi and 4dpi were then integrated as per the Seurat integration pipeline^10^ after normalizing the samples using SCTransform method. The first 30 dimensions were used with RunUMAP and FindNeighbors functions. For getting the subset of macrophages/monocytes, the largest cluster primarily comprised of macrophages/monocytes annotated by SingleR (BluePrintEncode database) was selected. Cells that were annotated as another cell type in this cluster were filtered out. The macrophages/monocytes from all BAL samples were then split into individual samples, normalized using SCTransform method and then integrated again using 30 dimensions.

The three lung samples from uninfected rhesus macaques were processed similarly. The following parameters were used to filter cells: (i) nFeature_RNA >=200 & <=4000, (iii) % of mitochondrial genes < 20, (iv) % of RPS/RPL genes < 50 and (v) log10(nFeature_RNA) / log10(nCount_RNA) >= 0.8. The samples were normalized using SCTransform and integrated. The first 40 dimensions were used for the initial clustering. The macrophage/monocyte cells as annotated by SingleR were then selected, split into individual samples and integrated again using 30 dimensions. Louvain clustering resulted in four clusters which were annotates based on the expression of marker genes. This integrated dataset served as the reference to map the macrophages/monocytes from SARS-CoV2 infected BAL using the FindTransferAnchors and MapQuery with reference.reduction set to pca and umap as the reduction.model. The BAL samples were also annotated using SingleR library with the IM and AM bulk sorted cells as reference.

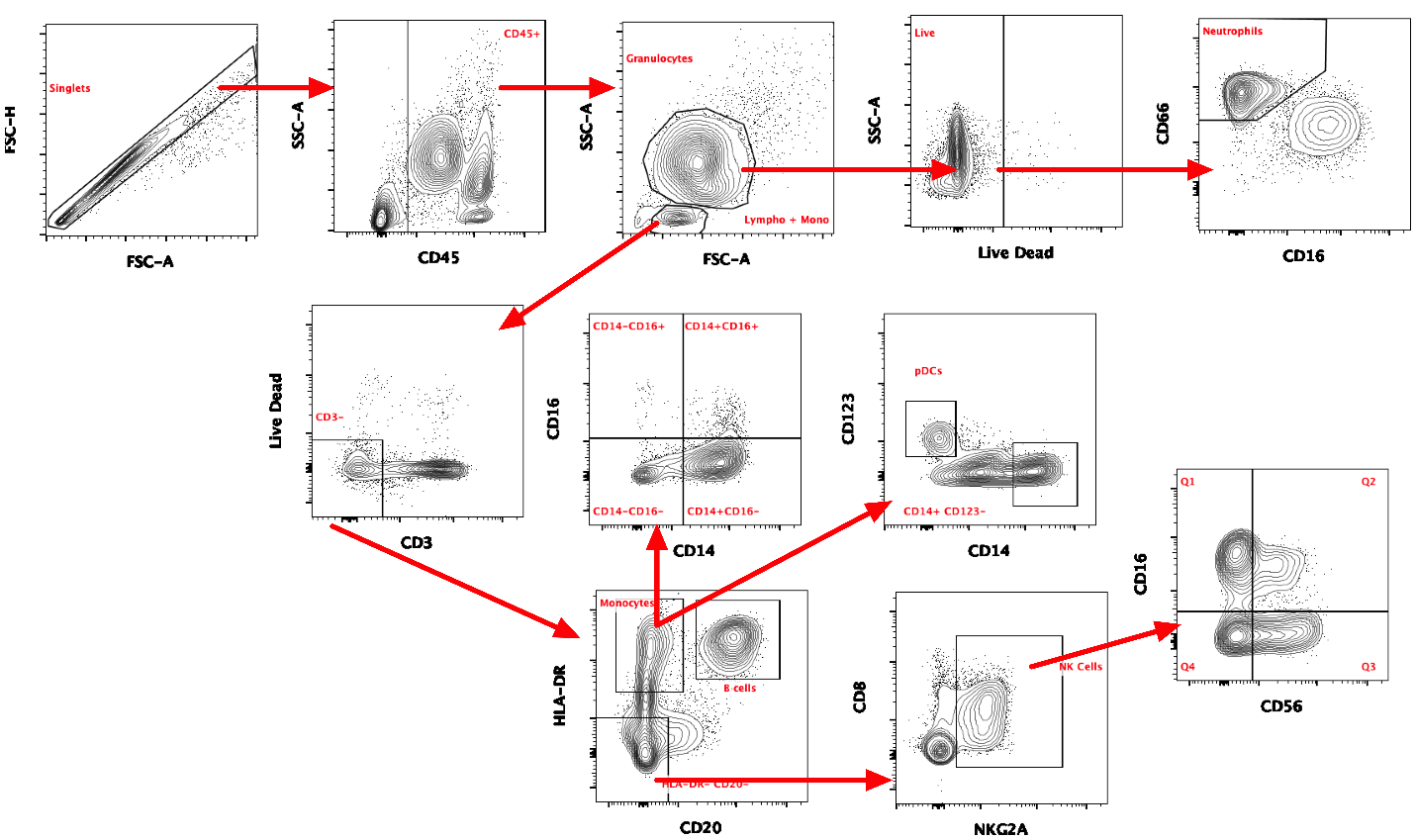

**Supplementary Figure 1. Flow sorting strategy for different immune cell populations.**

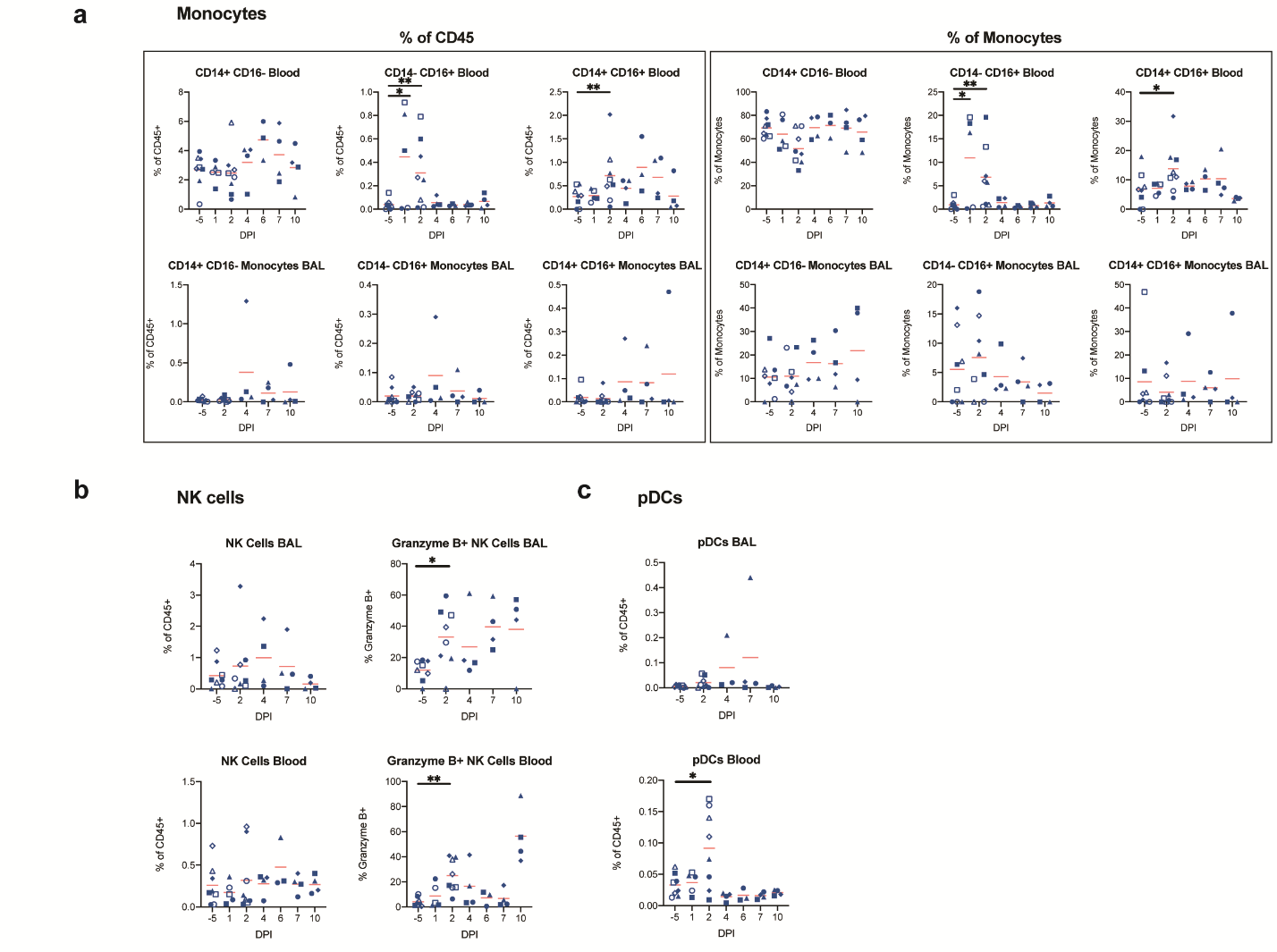

**Supplementary Figure 2. Longitudinal flow cytometric analysis in BAL and blood following SARS-CoV-2 infection.** (**A**) Longitudinal levels of monocytes within BAL and blood depicted as the percentage of CD45+ cells and % of monocytes (CD3^-^ CD20^-^ HLA-DR^+^). (**B**) Longitudinal levels of NK cells as a percentage of CD45+ cells and frequency of NK cells expressing Granzyme B in BAL and blood. (**C**) Longitudinal levels of plasmacytoid dendritic cells (pDCs) within BAL and blood depicted as a % of CD45+ cells. The red bars represent the mean. Statistical analysis was performed using one-tailed Wilcoxon signed-rank test comparing each timepoint to -5dpi. * p-value < 0.05, ** p-value < 0.01.

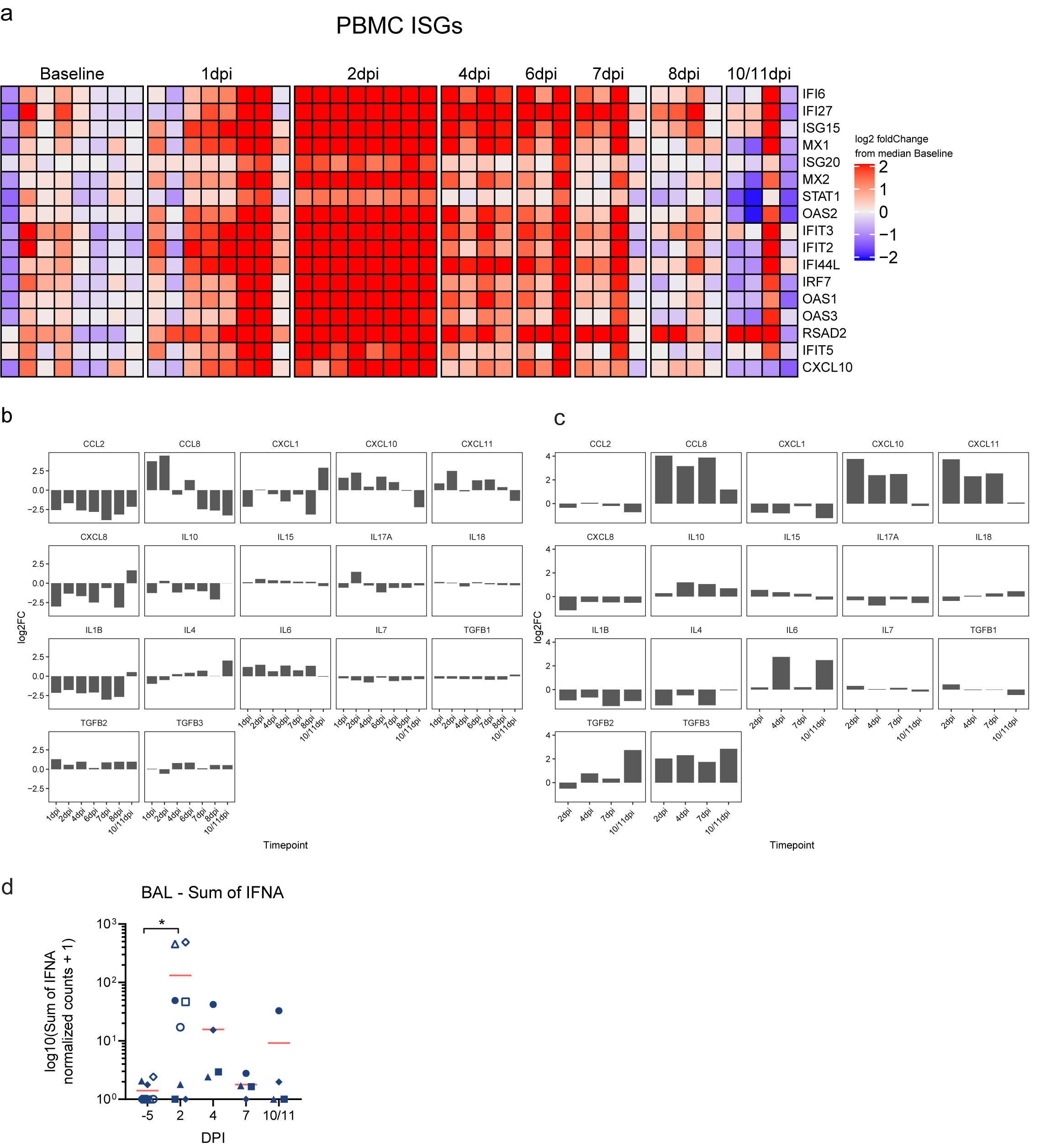

**Supplementary Figure 3. Bulk transcriptomic analysis of airways and peripheral blood. (A)** Heatmap showing expression of ISG in PBMC over all sampled time points. The color scale indicates log2 expression relative to the median of -5dpi samples. **(B & C)** Normalized expression of cytokines and chemokines in bulk RNA-Seq in BAL (B) and PBMC (C). **(D)** Sum of normalized expression of IFNA in longitudinal BAL samples. The red bars represent the mean. Statistical analysis was performed using one-tailed Wilcoxon signed-rank test comparing each timepoint to -5dpi. * p-value < 0.05.

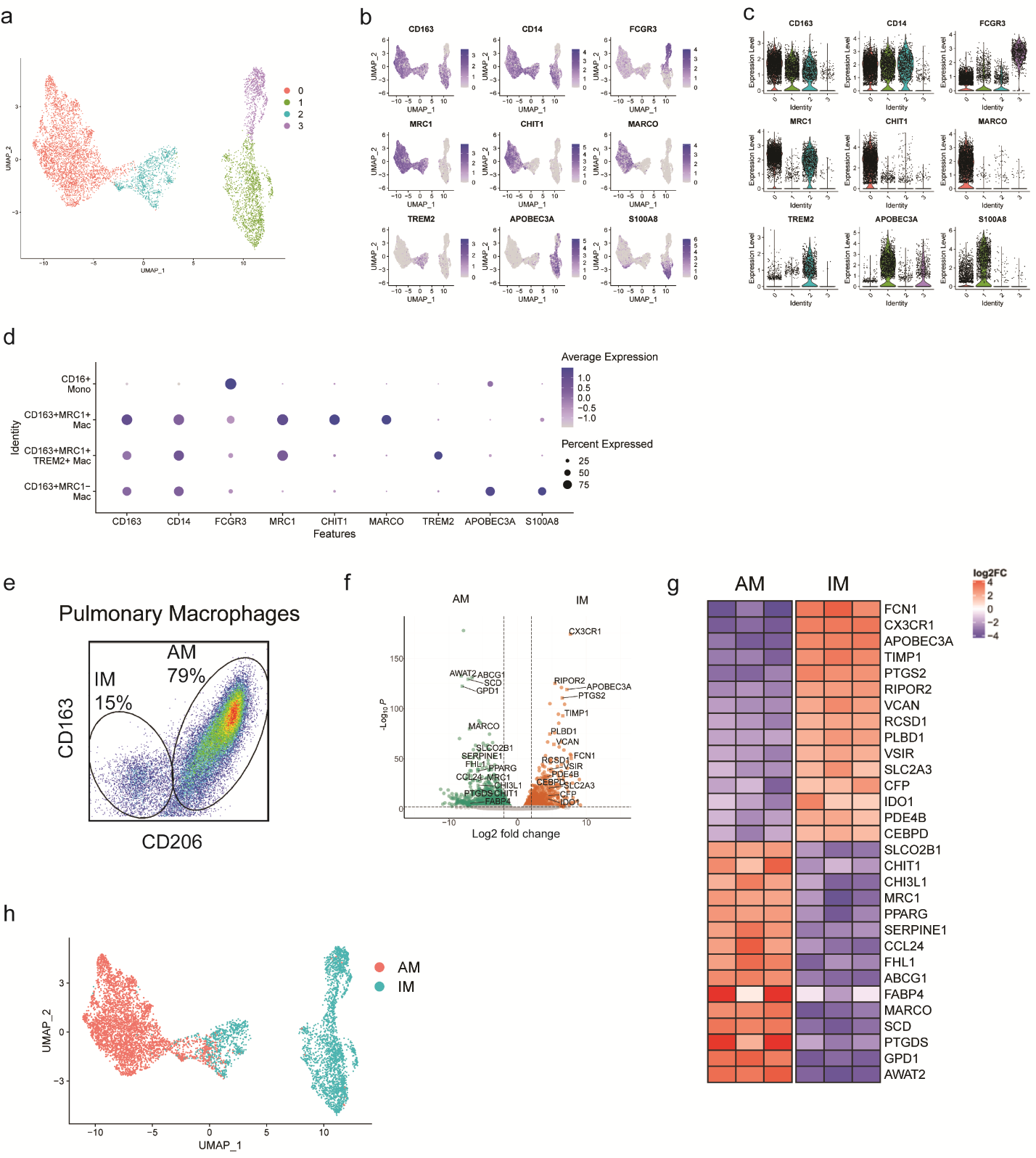

**Supplementary Figure 4. Reference used for annotating macrophage/monocyte subsets.**

**(a)** UMAP of macrophages/monocytes from 10X lung samples of three uninfected rhesus macaques (NCBI GEO: GSE149758) showing Louvain clustering. (**b & c**) FeaturePlot (b) and Violin Plots (c) showing the expression of marker genes in the macrophage/monocyte clusters. **(d)** DotPlot showing expression of marker genes for the different monocyte/macrophage subsets as defined previously ^11,12^ **(e)** Sorting strategy for interstitial and alveolar macrophages from lungs of three uninfected rhesus macaques **(f)** Volcano plots showing differentially expressed genes for pairwise comparison of alveolar and interstitial macrophages. The thresholds used are an adjusted p-value < 0.05 and a fold change of 2 for alveolar vs interstitial macrophage. Top 15 genes that have a mean normalized expression of at least 5000 for either type have been indicated. **(g)** Heatmap showing the top 15 genes for each subset. The color scale indicates log2 expression relative to the median of all samples. (**h**) UMAP of single-cell 10X lung samples showing SingleR annotations using the bulk sorted cells as reference.

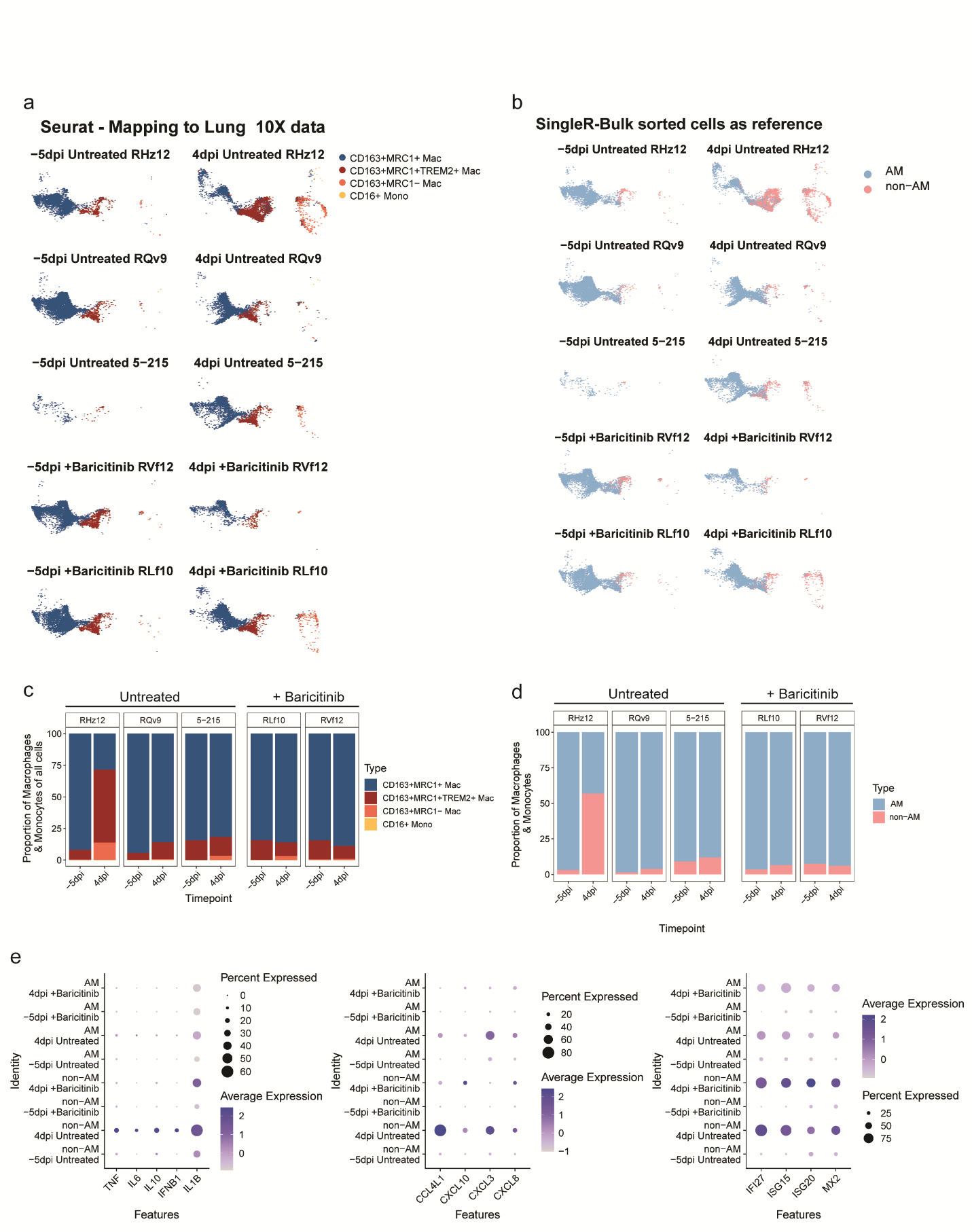

**Supplementary Figure 5. Cell annotation using bulk sorted cells as reference. (a & b)** The BAL macrophage/monocytes from SARS-CoV2 infected rhesus macaques (three untreated and two baricitinib treated) projected on the 10X lung reference macrophages/monocytes UMAP split by each animal and timepoint. (a) Annotations predicted from mapping to 10X lung samples using Seurat (b) Annotations predicted by SingleR using the bulk sorted AM and IM cells as reference.**(c & d)** Percentage of a macrophage/monocytes subset out of all the macrophage/monocyte cells in a given sample based on 10X lung reference (c) and the bulk sorted cells as reference (d). **(e)** DotPlots showing expression of pro-inflammatory cytokines, chemokines and ISG in macrophage/monocyte subsets based on the bulk sorted cells reference at -5dpi and 4dpi in BAL samples from untreated and baricitinib treated rhesus macaques.

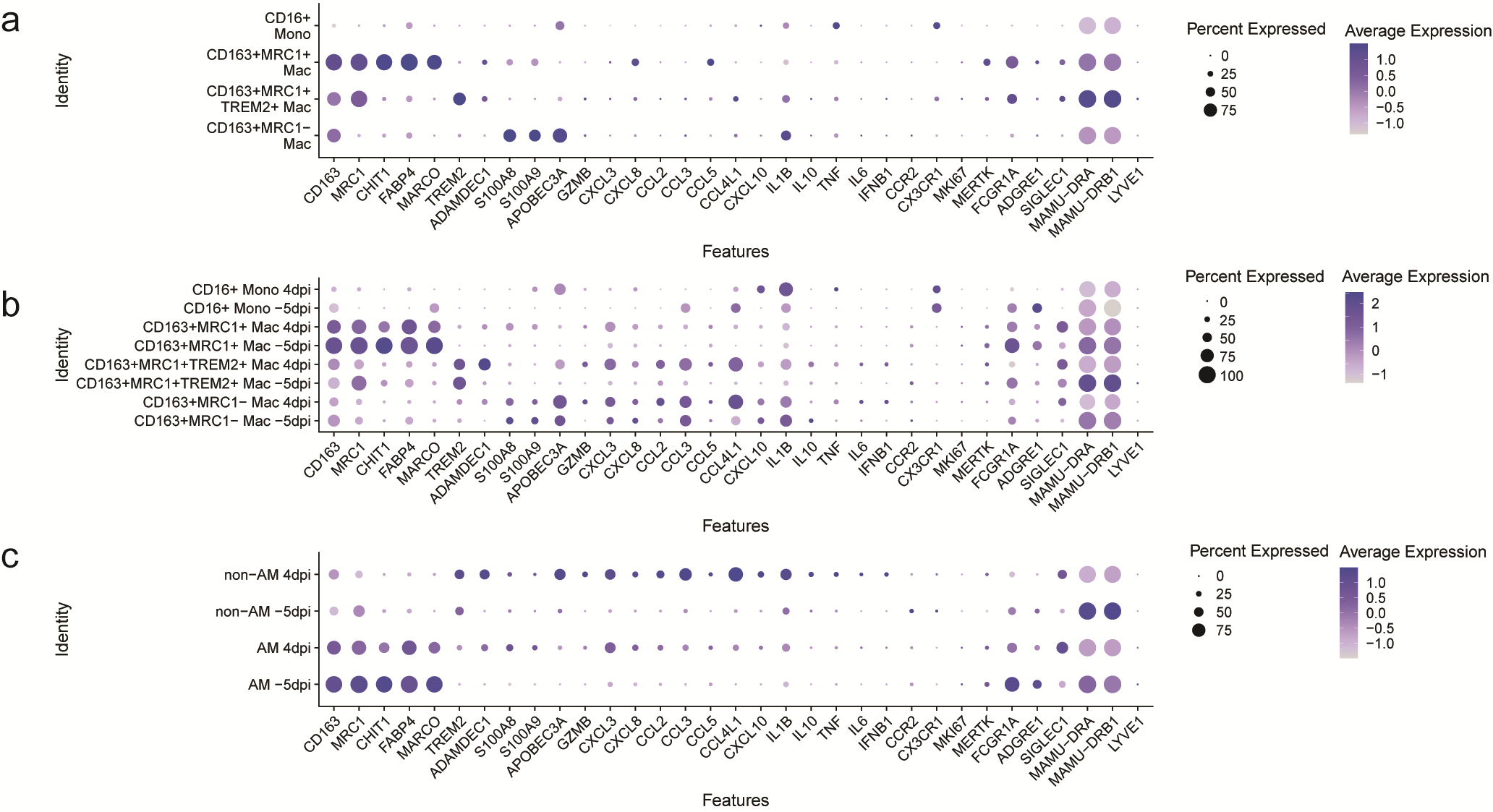

**Supplementary Figure 6.** Expression of DE genes in different macrophage/monocyte subsets in **(a)** 10X lung control samples from three uninfected RM. **(b & c)** BAL samples from three SARS-CoV2 infected RM annotated using the 10X lung reference (b) or the bulk sorted cells as reference (c).
